# Polyserine-tau interactions modulate tau fibrillization

**DOI:** 10.1101/2025.06.03.657713

**Authors:** James Pratt, Kathleen McCann, Jeff Kuo, Roy Parker

## Abstract

Tau aggregates are the defining feature of multiple neurodegenerative diseases and contribute to the pathology of disease. However, the molecules affecting tau aggregation in cells are unclear. We previously determined that polyserine-rich domain containing proteins enrich in tau aggregates, assemble into puncta that can serve as sites of tau aggregation, and exacerbate tau aggregation in cells and mice. Herein, we show that polyserine domains are sufficient to define assemblies as sites of tau aggregation, in part, through localization of tau seeds. Purified polyserine self-assembles and directly interacts with monomeric and fibrillar tau. Moreover, polyserine-tau assemblies recruit RNA, leading to faster rates of tau fibrillization *in vitro*. Using polyserine variants, we found that enrichment in tau aggregates and stimulation of tau aggregation are separable functions of polyserine domains, with polyserine self-assembly stimulating tau aggregation. Together, our results show that polyserine self-assembles and directly interacts with tau to form preferred sites of tau aggregation.

## Introduction

The aggregation of tau is a hallmark of several neurodegenerative diseases, including Alzheimer’s Disease (AD) (1). Supporting a causal role for tau in disease, symptom severity in AD correlates with increased tau pathology (2). Importantly, both genetic and environmental inputs can promote tau aggregation. For example, genetic mutations in *MAPT* can result in coding variants with an increased propensity for aggregation, as in frontotemporal dementia (FTD) (3). In contrast, secondary triggering events such as repetitive head trauma or amyloid-β plaques, can increase tau aggregation in chronic traumatic encephalopathy (CTE) (4), or AD (5), respectively. While tau plays an important role in disease contexts, the intracellular mechanisms of tau aggregation, and the impact of additional proteins or other cofactors on tau aggregation, are generally unknown.

We and others have observed that tau aggregates contain other molecules, including RNA and specific RNA binding proteins (6–10), which may affect tau aggregation in cells. Previously, we observed that tau aggregates enrich small structured RNAs, such as snRNAs and snoRNAs (6). Additionally, the splicing speckle proteins Serine-Argine Repetitive Matrix 2 (SRRM2) and Pinin (PNN) mislocalize to cytoplasmic tau aggregates in cell culture models, PS19 tau transgenic mice, and in human disease (6, 9). SRRM2 and PNN are normally present in nuclear speckles that colocalize with nuclear tau aggregates in cell line and mouse models (6). Additionally, SRRM2 and PNN can mislocalize into cytoplasmic tau aggregates through formation of cytoplasmic speckles (CSs), or mitotic interchromatin granules (MIGs) that act as preferred sites of tau aggregation (7).

The ability of SRRM2 and PNN to interact with, and enhance the formation of tau aggregates is based on their unique polyserine domains (7). SRRM2 contains two C-terminal polyserine domains of 25 or 42 residues, while PNN contains a C-terminal serine rich domain with 50 of 66 residues being serine (7). These polyserine domains are both necessary and sufficient to target exogenous proteins to tau aggregates (7). In addition, three observations demonstrate these domains can affect the efficiency of tau aggregation. First, knockdown of PNN results in decreased tau aggregation in cells (7). Second, overexpression of the SRRM2 or PNN C-terminal domains, or the 42 amino acid long polyserine domain (polyserine_42_), is sufficient to increase tau aggregation in HEK293T tau biosensor cells (7). Finally, polyserine_42_ expression in PS19 mice exacerbates tau pathology based on increased phosphorylated tau and increased seeding capacity of brain homogenate (11). It remains unclear whether polyserine itself self-assembles to form intracellular assemblies, if there is a direct interaction between tau and polyserine domains, or if polyserine domains are sufficient to convert other subcellular assemblies into preferred sites of tau aggregation.

To examine the physical interaction between polyserine domains and tau, and how those interactions affect the process of tau aggregation, we performed a series of *in cellulo* and *in vitro* experiments. We demonstrate that stress granules, a cytoplasmic assembly of RNA and proteins (12), can be converted to a preferred site of tau aggregation through expression of a G3BP1-polyserine_42_ fusion protein. We next used *in vitro* microscopy and kinetic experiments to characterize the tau-polyserine domain interaction. First, we demonstrate purified polyserine_42_ forms self-assemblies. Second, we demonstrate polyserine_42_ assemblies directly interact with tau monomers and seeds and can increase the rate of tau fiber formation and growth. The surface of the resulting tau fibers is decorated with polyserine assemblies. Finally, we show that the self-assembly of polyserine_42_ is separable from enrichment in tau aggregates, and self-assembly correlates with effects on tau aggregation. Taken together, these observations demonstrate polyserine domains form self-assemblies that can directly interact with tau to promote aggregation.

## Results

### Polyserine_42_ Converts Stress Granules into Sites of Tau Aggregation

We and others have previously observed that G3BP1 positive stress granules are not sites of tau aggregation (7, 13). In contrast, assemblies of overexpressed polyserine_42_ and cytoplasmic speckles containing polyserine rich proteins, serve as preferred sites of aggregation in tau biosensor cells (7). The difference in tau aggregation between stress granules and polyserine containing assemblies could be because polyserine domains are necessary and sufficient to define a preferred site of tau aggregation, or because stress granules contain inhibitors of tau aggregation. This latter possibility is suggested by the observations that the stress granule component G3BP2 can limit tau aggregation *in vitro* and in neurons (14). To distinguish these possibilities, we examined if polyserine_42_ could convert stress granules into sites of tau aggregation.

To do so, we constructed tau biosensor cell lines stably expressing either mRuby, mRuby-G3BP1^10^, or mRuby-G3BP1-polyserine_42_ (Fig 1A). The HEK293T tau biosensor cells express cyan fluorescent protein (CFP) or yellow fluorescent protein (YFP) tagged forms of the microtubule binding domain of tau with a P301S pathogenic mutation that forms FRET (Förster Resonance Energy Transfer) positive tau aggregates after transfection of exogenous tau seeds (15). The proper expression of these G3BP1 fusion proteins was verified by western blot (Fig S1A). After seeding with brain extracts from Tg2541 mice, cells were treated with the translational inhibitor pateamine A (PatA) to induce irreversible stress granule formation (16). This experiment provided several notable observations.

**Figure 1:**
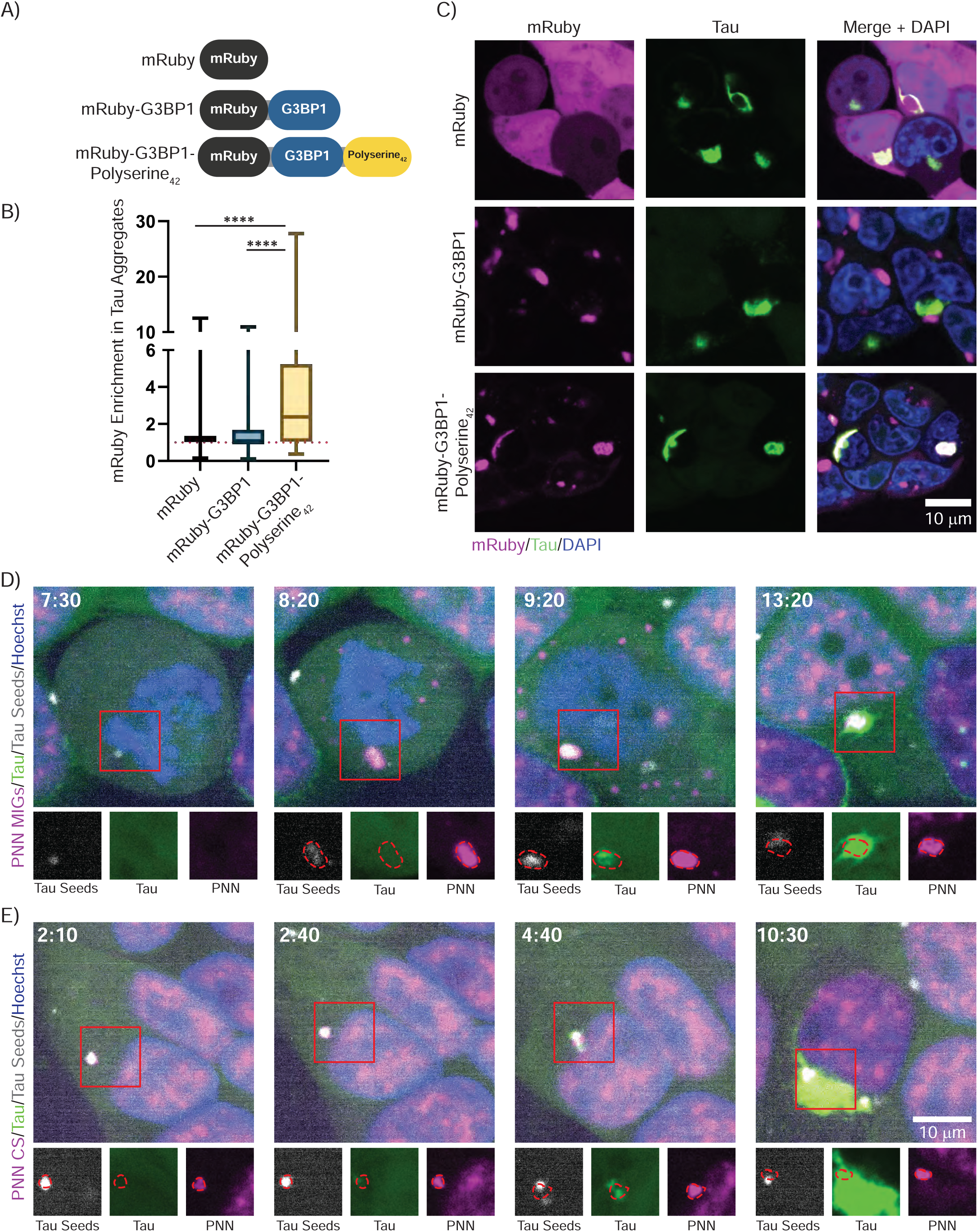
Polyserine_42_ Containing Assemblies Are Sites of Tau Aggregation Through Enrichment of Tau Seeds. (A) Schematic of mRuby fusion proteins stably expressed in HEK293T tau biosensor cells. (B) Microscopy of HEK293T tau biosensor cells stably expressing mRuby, mRuby-G3BP1, or mRuby-G3BP1-Polyserine_42_ (magenta) were transfected with brain homogenate from aged Tg2541 mice to seed tau aggregates (green) and fixed twenty-four hours after transfection, nuclei stained with DAPI (blue). (C) Quantification of mRuby enrichment in cytoplasmic tau aggregates. N = at least 164 cells from 3 biological replicates. Statistics performed with Kruskal-Wallis test with Dunn’s multiple comparisons test. (****) P < 0.0001 (D) Stills from live cell imaging of HEK293T tau biosensor cells with endogenously tagged PNN-Halo (magenta) MIGs, transfected with fluorescent tau seeds (grey), to induce endogenous tau aggregation (green), nuclei stained with DAPI (blue). Time in hours:minutes post transfection and start of imaging. Channel breakouts from each time point in greyscale below with PNN assemblies outlined in red and superimposed to each channel. (E) Live cell imaging as performed in (D) of PNN positive CS.

We observed mRuby-G3BP1 and mRuby-G3BP1-polyserine_42_ assemblies form in the cytoplasm of ∼10% of cells in the absence of PatA treatment and tau seeding (Fig S1B-D). Since PolyA Binding Protein (PABP), another stress granule marker, did not localize to these assemblies these are presumably polyserine-driven assemblies of mRuby-G3BP1-polyserine_42_ protein (Fig S1C). Treatment with PatA induced mRuby positive assemblies which enriched PABP in greater than 30% of cells in either G3BP1 or G3BP1-polyserine_42_ expressing cell lines (Fig S1C and D). Therefore, the addition of polyserine_42_ to G3BP1 does not block the formation of canonical stress granules. We did not observe significant differences in the amount of tau aggregation after seeding with and without treatment with PatA across the three cell lines (Fig S1E). Therefore, the formation of stress granules by translation inhibition does not change tau aggregation in cells.

A key result was that stress granules containing G3BP1-polyserine_42_ colocalized strongly with cytoplasmic tau aggregates in PatA treated and seeded cells (Fig 1B and C). In contrast, we observed that tau aggregates and G3BP1 positive stress granules without polyserine_42_ did not colocalize (Fig 1B and C), as previously reported (7, 13). This demonstrates that targeting polyserine domains to stress granules allows those organelles to interact with tau aggregates.

To understand if G3BP1-polyserine_42_ stress granules act as sites of tau aggregation, or merge with tau aggregates after their formation, we performed live cell imaging. We found that G3BP1 or G3BP1-polyserine_42_ positive stress granules formed before tau aggregation began (Movie S1 and S2, Fig S2A). In cells with G3BP1 positive stress granules, tau aggregation initiated separately from stress granules (Fig S2A, top panels). In cells expressing G3BP1-polyserine_42_, tau aggregation initiated within the stress granule and remained colocalized with G3BP1-polyserine_42_ throughout imaging (Fig S2A, bottom panels). Thus, stress granules that contain polyserine_42_ can serve as sites of tau aggregation initiation.

Taken together, this indicates that the presence of polyserine_42_ is sufficient for converting stress granules into a preferred site of tau aggregation. This also demonstrates that stress granules do not contain endogenous inhibitors of tau aggregation.

### Polyserine containing assemblies concentrate tau seeds

One possible mechanism by which assemblies containing polyserine domains could act as sites of tau aggregation would be to recruit nascent tau seeds as they enter the cell and then serve as a site for those seeds to grow. To examine if tau seeds interacted with polyserine domain containing assemblies, we transfected tau seeds prepared *in vitro* from fluorescently labeled tau and followed the fate of those seeds in tau biosensor cell lines expressing SRRM2-Halo or PNN-Halo (7).

Strikingly, we observed that transfected tau seeds localized within SRRM2 positive MIGs and CSs (Movies S3 and S4, Fig S3A and B), and PNN positive MIGs and CSs, which then led to nucleation of tau aggregate growth (Movies S5 and S6, Fig 1D and E). Similarly, we observed fluorescent tau seeds localized in stress granules with G3BP1-polyserine_42_ (Movie S7, Fig S2B) but not in wild-type stress granules (Movie S8, Fig S2C). These observations support that polyserine containing assemblies serve as sites of tau aggregation, at least in part, by localizing tau seeds.

### Polyserine_42_ Self-Assembles *In Vitro*

Our previous observations that polyserine_42_ overexpression forms cytoplasmic assemblies (7) demonstrated that polyserine domains can promote the assembly of non-membrane bound organelles in cells. To determine if this was driven by possible self-assembly properties of polyserine, we purified and examined the properties of a 42 amino acid long polyserine domain, as found in the C-terminus of SRRM2 (7). We purified polyserine_42_ with a TEV-cleavable, N-terminal SUMO tag for solubility, and a C-terminal Halo tag to facilitate visualization. As a control, we purified Halo with a TEV-cleavable, N-terminal SUMO tag (Fig 2A). To monitor assembly by confocal microscopy we labeled polyserine_42_-Halo and Halo with a Janelia Fluor 646-Halo ligand.

**Figure 2:**
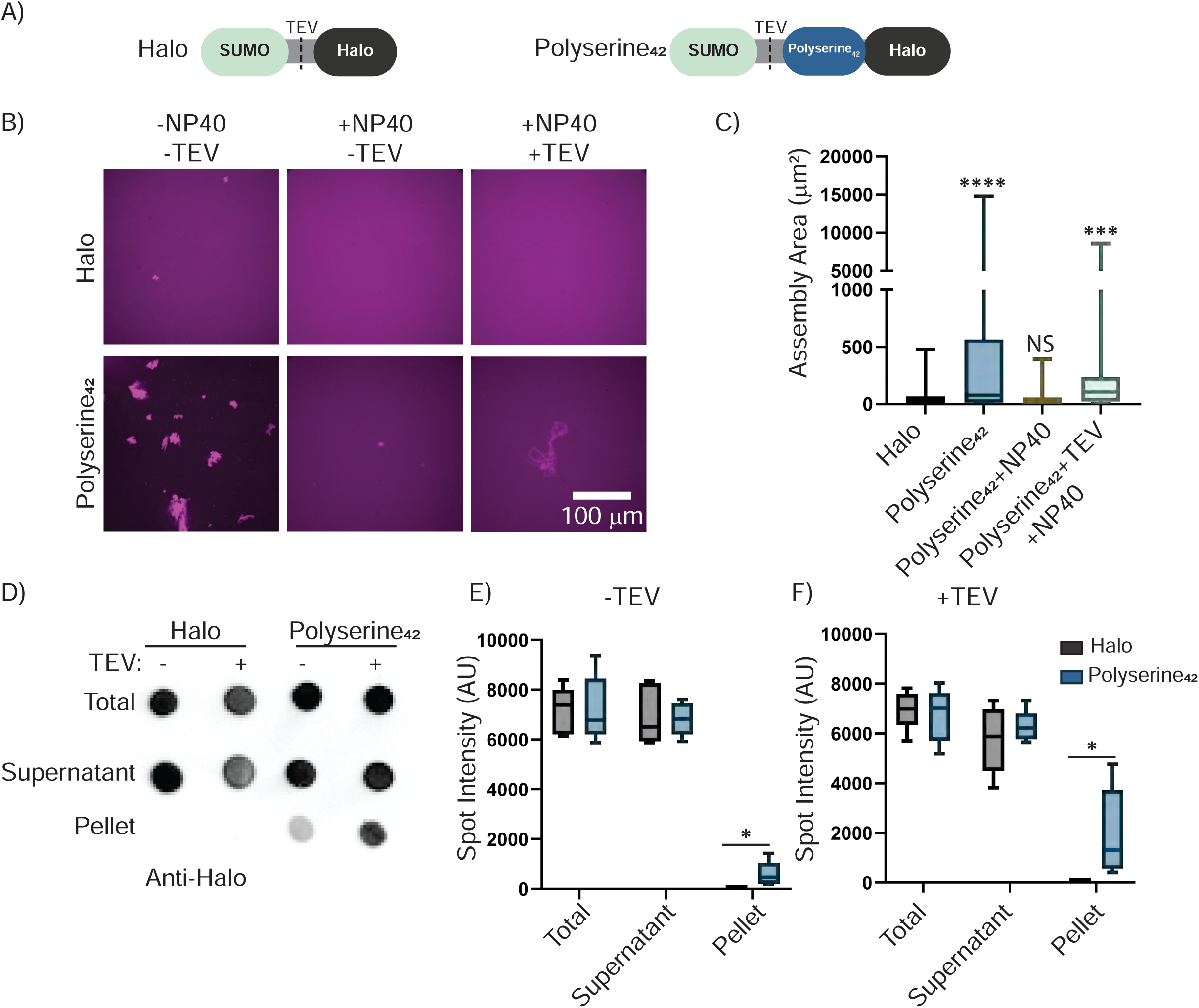
Polyserine_42_ Self-Assembles *in vitro*. (A) Schematic of recombinant SUMO(TEV)-Halo and SUMO(TEV)Polyserine_42_-Halo constructs. (B) Representative images of Halo or Polyserine_42_-Halo labeled with Janelia-Fluor 646 Halo Ligand purified without detergent or TEV protease treatment (left), with 0.1% NP40 (middle), and with both 0.1% NP40 and TEV protease treatment (right). (C) Quantification of assembly size. N= at least 35 assemblies across 3 biological replicates, Statistics were performed with Kruskal-Wallis with Dunn’s Multiple Comparisons test. (***) P = 0.006 (****) P < 0.0001. (D) Representative dot blot of Halo or Polyserine_42_-Halo stained with Anti-Halo from ultracentrifugation pelleting experiments performed in 0.1% NP40 with and without TEV protease treatment. (E) Quantification of dot intensity from N = 3 biological replicates of dot blot as in (D) without TEV treatment. Statistics performed with Mann-Whitney U test across Halo (Black) or Polyserine_42_-Halo (Blue) from Total, Supernatant, or Pellet fractions (*) P < 0.005 (F) Quantification of dot intensity from N = 3 biological replicates of dot blot as in (D) with TEV treatment. Statistics performed with Mann-Whitney U test across Halo (Black) or Polyserine_42_-Halo (Blue) from Total, Supernatant, or Pellet fractions (*) P < 0.005.

Following purification, we observed polyserine_42_ rapidly and spontaneously self-assembles *in vitro*. Polyserine_42_ forms large, non-spherical assemblies, while the halo control only formed rare, much smaller, assemblies (Fig 2B and C). When purified in the presence of 0.1% NP-40, polyserine_42_ still forms small assemblies, suggesting they are partially resistant to solubilization with detergent (Fig 2B and C). A fraction of purified polyserine_42_ pellets when subjected to ultracentrifugation, providing further evidence for polyserine_42_ assembly (Fig 2D-F). Removal of the SUMO solubility tag through TEV cleavage increases the size of polyserine_42_ assemblies purified in detergent, and the amount of pelleted polyserine_42_ (Fig 2B-F). Thus, self-assembly is an intrinsic feature of polyserine domains.

### Polyserine_42_ assemblies directly interact with monomeric and seed competent tau

Polyserine_42_ assemblies in cells can be sites of tau aggregation (7). Moreover, we observed that tau seeds could be localized to polyserine containing assemblies such as MIGs, CSs, and stress granules tagged with polyserine_42_. This suggests two possible manners by which polyserine_42_ might interact with tau and affect aggregation. First, polyserine_42_ might form a structure and/or organization that specifically binds tau as a monomer and/or fiber. Second, polyserine_42_ assemblies could bind proteins promiscuously, through a network of hydrogen bond donors and acceptors. To understand if polyserine_42_ assemblies are directly interacting with tau or tau seeds, we asked if tau monomers or tau seeds could associate with self-assembled polyserine_42_.

We purified full length 2N4R P301S tau with a single cysteine to allow for direct observation of tau via maleimide labeling with fluorescent dyes (Fig 3A). We mixed monomeric tau with polyserine_42_-Halo assemblies or Halo alone and performed confocal microscopy to monitor potential tau and polyserine_42_ interactions. Tau does not alter the polyserine_42_ self-assembly in vitro (Fig S4A) and is not required for polyserine_42_ assembly in WT HEK293T cells (Fig S4B).

**Figure 3:**
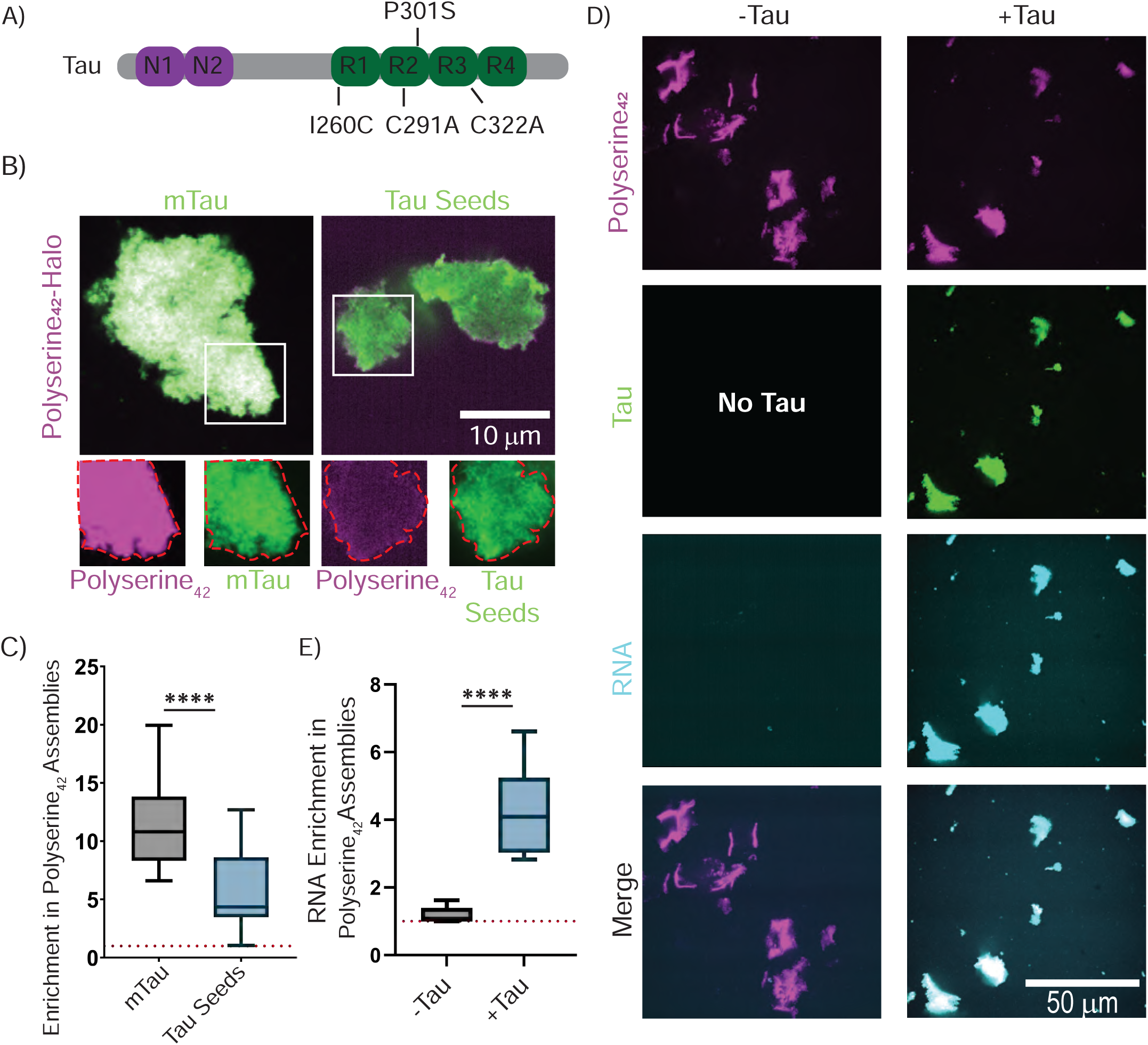
Polyserine_42_ Interacts Directly with Tau Monomers and Seeds. (A) Schematic of recombinant tau construct used. (B) Representative images of tau monomers (mTau) maleimide labeled with JF-646, tau seeds prepared from mTau maleimide labeled with JF-646 incubated with shaking with polyU RNA (green) mixed with Polyserine_42_-Halo labeled with JF-549. Channel breakouts shown below each image in grayscale with polyserine assembly perimeter in red superimposed to other channels. (C) Quantification of protein enrichment in Polyserine_42_-Halo assemblies as in (B). Statistics performed with one-way Kruskal-Wallis test with Dunn’s multiple comparisons test. (***) P < 0.005 (****) P < 0.0001. (D) Representative images of Polyserine_42_-Halo assemblies (magenta) incubated with fluorescent polyU RNA (cyan) with and without tau monomers (green). Channel breakouts in grayscale. (E) Quantification of RNA enrichment in Polyserine_42_-Halo assemblies with and without tau monomers as in (D). Statistics performed with Mann-Whitney U test. (****) P < 0.0001

Importantly, we observe that tau monomers enrich in polyserine_42_-Halo assemblies suggesting a direct interaction (Fig 3B and C). To rule out effects from the SUMO or Halo tags, we also purified polyserine_42_ with a C-terminal cysteine to allow for maleimide labeling, and a TEV-cleavable N-terminal SUMO tag for solubility (Figure S4C) We observe polyserine_42_-Cys assemblies also interact with tau monomers directly (Fig S4 D and E). Interestingly, polyserine_42_-cys assemblies did not interact with GFP or α-synuclein (Fig S4D and E). Therefore, polyserine_42_ directly interacts with tau monomers with some degree of specificity.

Since we observed tau seeds interacting with polyserine domain containing assemblies in cells (Fig 1C), we also examined if tau seeds prepared by sonication of fluorescently labeled tau fibers enriched in polyserine_42_ assemblies *in vitro*. We observed tau seeds could associate with polyserine_42_-Halo and polyserine_42_-Cys assemblies, although to a lesser degree than tau monomers (Fig 3B and C, S4D and E). This suggests that the recruitment of tau seeds into polyserine-containing assemblies in cells could occur through direct interaction.

We also examined if polyserine_42_ assemblies enrich RNA, a co-factor sufficient for inducing tau fiber formation (17). We mixed polyserine_42_-Halo assemblies with fluorescently labeled polyU RNA. We observed polyserine_42_-Halo assemblies do not enrich polyU RNA (Fig 3D and E). However, in the presence of tau we observed that tau, RNA, and polyserine_42_-Halo co-assemble (Fig 3D and E). Therefore, polyserine_42_ assemblies enrich RNA in a tau dependent manner.

### Polyserine_42_ Increases the Rate of Tau Fiber Formation and Growth

Since polyserine_42_ can interact with tau, and thereby potentially either alter the folding of tau and/or create a high local concentration of both tau and RNA, it may directly increase rates of tau aggregation. To measure the rate of tau fiber formation and growth we used an *in vitro* thioflavin-t (ThT) kinetics assay. First, we formed tau fibers with polyU RNA and increasing concentrations of tau in the presence of ThT to determine the optimal concentration to see changes in fibrilization rates and lag times (Fig S5A-C). The resulting ThT fluorescence curves were fit to a logistic growth equation, and lag times and rate constants of fibrilization were determined (Fig S5A-C). We found that tau fibrilization at 5 µM was optimal to see changes in rate constants and lag times.

We next performed the same ThT kinetics assay with 5 µM tau, polyU RNA, with or without polyserine_42_ or halo. We mixed monomeric 2N4R P301S tau with polyU RNA, which is sufficient to induce tau fibrillization (Fig 4A yellow curve), with polyserine_42_ and polyU RNA (Fig 4A black curve) or halo and polyU RNA (Fig 4A blue curve) and measured the increase in bound ThT fluorescence over time.

**Figure 4:**
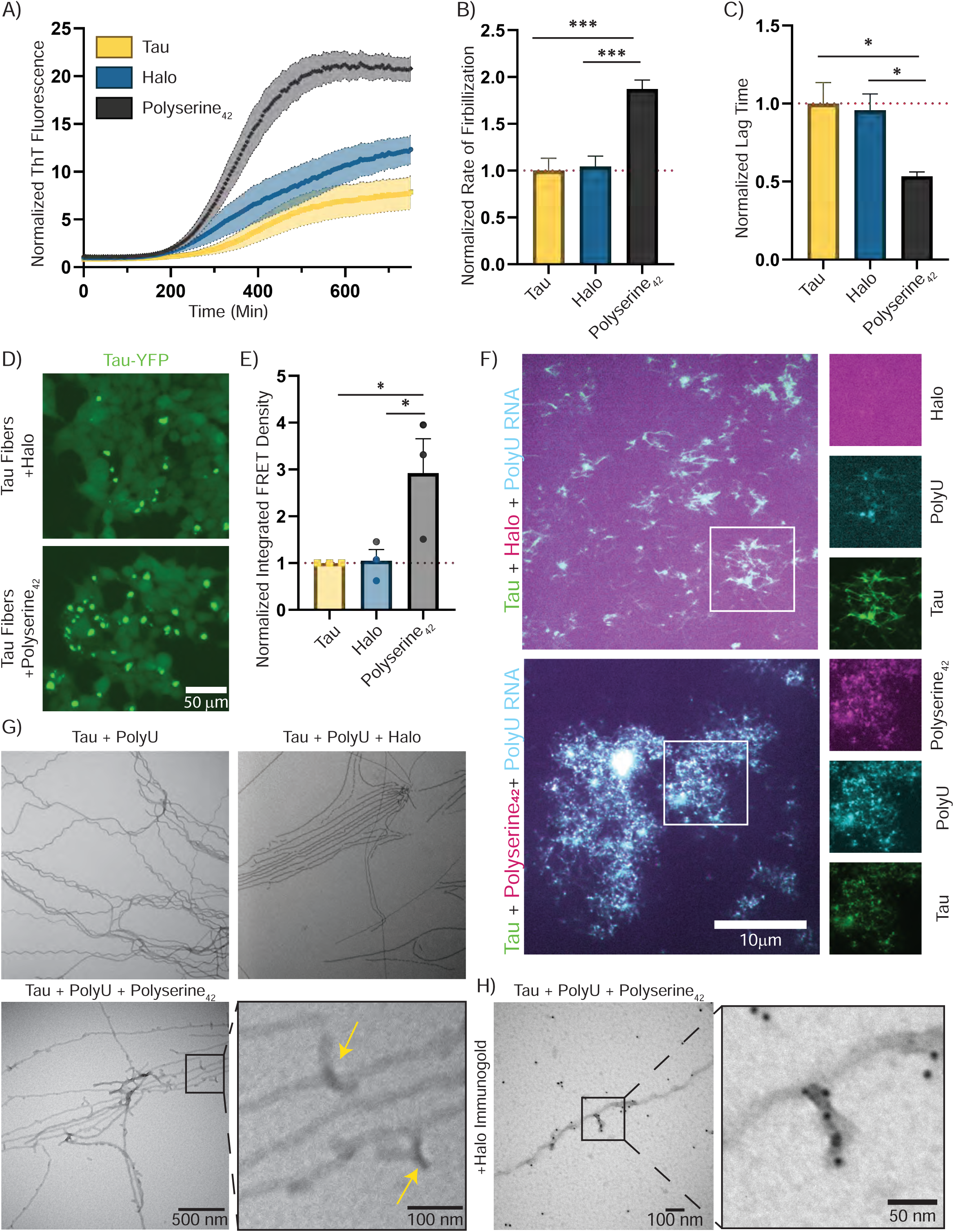
Polyserine_42_ Increases the Rate of Tau Fiber Formation and Alters Tau Fiber Ultrastructure. (A) Monomeric tau was incubated with PolyU RNA, with and without Halo or Polyserine_42_-Halo and thioflavin-t (ThT). ThT fluorescence was measured every 5 minutes for 24 hours, X-axis truncated to show first 800 minutes. Fluorescence normalized to first time point across each condition. N = 9 technical replicates across 3 biological replicates. (B) Quantification of the rate constants from logistic growth fitting of curves shown in (A). N = 9 technical replicates across 3 biological replicates. Statistics performed with Kruskal-Wallis test. (*) P < 0.05 (***) P < 0.001 (C) Quantification of the lag time from logarithmic growth fitting of curves shown in (A). Statistics performed with Kruskal-Wallis test. (*) P < 0.05 (**) P< 0.005, (***) P < 0.001 (D) Representative images of HEK293T tau biosensor cells transfected with tau fibers prepared with monomeric tau, polyU RNA, and Halo (top) or with Polyserine_42_-Halo (bottom) to seed tau aggregates (green). (E) Quantification of integrated FRET density (product of FRET+ percentage and median fluorescence intensity) of HEK293T tau biosensor cells transfected with tau fibers prepared with polyU RNA with and without Halo or Polyserine_42_-Halo normalized to cells transfected with tau fibers prepared without Halo or Polyserine_42_-Halo). N = 3 biological replicates. Statistics performed with Kruskal-Wallis test. (*) P < 0.05. (F) Representative images of tau fibers prepared with tau monomers (green), FITC-fluorescent polyU RNA (cyan), and Polyserine_42_-Halo or Halo labeled with JF646. Channel breakouts and zooms shown in gray scale. (G) Representative negative stain electron microscopy images of tau fibers prepared as in F. Yellow arrows show additional densities present in Polyserine_42_-Halo containing tau fiber preps. (H) Representative image of immunogold negative stain electron microscopy of tau fibers prepared as in F stained with anti-halo and secondary antibody conjugated with gold nanoparticles, zoom and inset shows a Polyserine_42_-Halo assembly on the surface of a tau fiber.

Importantly, we observed that polyserine_42_ significantly decreases the lag time and increases the rate constant of fibrilization compared to halo or tau alone (Fig 4A-C black bars). To test if polyserine_42_ was increasing the rate of tau fibrilization by acting as an additional cofactor, we performed the same ThT assay with and without polyserine_42_ or halo in the absence of polyU RNA. We observed that polyserine_42_ nor halo are sufficient cofactors for tau fiber formation (Figure S5D). This suggests that polyserine_42_ can increase tau fibrillization, potentially by increasing the rate of formation of initial seeding competent tau species, and the rate of growth of those seeds.

To examine if polyserine_42_ was increasing the conversion of tau into fibrillar species by an orthogonal assay, we asked if polyserine_42_ increases the amount of seeding competent tau following *in vitro* fibrillization. In this experiment, we transfected tau fibrilization reactions after 24 hours of incubation with or without polyserine_42_, or Halo into HEK293T tau biosensor cells (15). In agreement with the increased rate of tau fiber formation and growth, we found that the presence of polyserine_42_ results in an increase in the amount of seeding competent tau as compared to Halo alone (Fig 4D and E). This provides additional evidence that polyserine_42_ increases the formation of fibrillar, seeding competent tau.

### Polyserine_42_ Incorporates into Growing Tau Aggregates and Changes Tau Fiber Ultrastructure

After observing that polyserine_42_ directly increases the rate of formation and growth of tau fibers, we asked if the resulting fiber structures and composition are altered. Confocal and negative stain electron microscopy were performed to determine the composition and ultrastructure, respectively, of tau fibers. We prepared tau fibers as described previously and imaged fibers after 24 hours of fibrilization.

With confocal microscopy we found that tau fibers prepared in the presence of polyserine_42_ show colocalization of polyserine_42_, tau, and polyU RNA (Fig 4F). Tau fibers prepared in the presence of halo alone show diffuse halo signal, while tau and polyU RNA fluorescence is colocalized (Fig 4F).

Negative stain electron microscopy of tau-RNA fibers showed polyserine_42_ can change tau fiber ultrastructure. Tau fibers made in the presence of polyserine_42_ contain additional densities decorating the fiber (Fig 4G, yellow arrows) which are not observed with tau fibers prepared with Halo, or no additional protein. To identify these unique densities, we performed immunogold labeling against the Halo tag. We observed the enrichment of gold-nanoparticle densities on the extra-densities observed with tau-polyserine_42_ fibers (Fig 4H). We did not observe staining throughout the tau fiber, indicating that polyserine_42_ may interact with the surface of fibers, as opposed to incorporating into growing tau fibers. Taken together, this suggests that polyserine_42_ can interact with and possibly alter the form of growing tau fibers.

### Polyserine-tau interaction is separable from increasing tau aggregation

The results above show polyserine_42_ self-assembles, localizes to tau aggregates, and increases tau fibrillization. In principle, these effects could all be due to one biochemical property of polyserine_42_ or might reflect different functions. To address the possible overlap of these functions, we created variants of polyserine_42_ and examined their ability to localize to tau aggregates, affect the degree of tau aggregation, and form intracellular assemblies in HEK293T tau biosensor (15) cells. We created variants where a fraction of serine residues was substituted with negatively charged aspartic acid, positively charged lysine, or neutral threonine residues without altering peptide length (Figure 5A).

**Figure 5:**
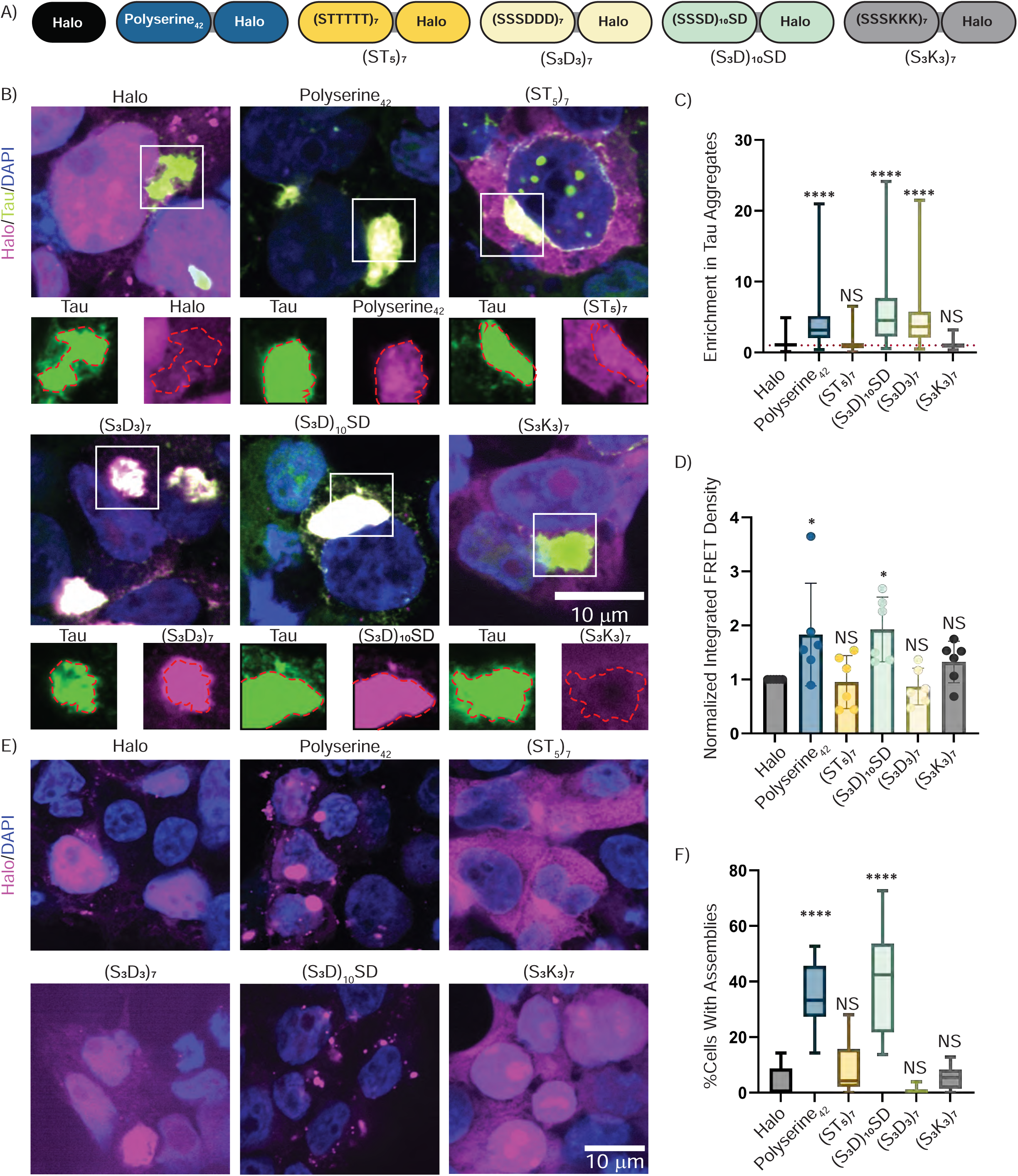
Analysis of Polyserine_42_ Variants. (A) Schematic of Polyserine_42_-Halo variants used. (B) Representative images of HEK293T tau biosensor cells transfected with plasmids expressing Halo, Polyserine_42_-Halo, or polyserine_42_-Halo variants labeled with JF646 halo-ligand (magenta) for 24 hours, then transfected with tau brain homogenate from aged Tg2541 mice to induce endogenous tau aggregation (green), and fixed after 24 hours, nuclei stained with DAPI (blue). Channel breakouts show tau aggregate perimeter in red superimposed to Halo/ Polyserine_42_-Halo /Polyserine_42_-Halo variant channels. (C) Quantification of Halo/ Polyserine_42_-Halo /Polyserine_42_-Halo variants enrichment in cytoplasmic tau aggregates. N = at least 129 cells with tau aggregates and Halo/ Polyserine_42_-Halo/Polyserine_42_-Halo variant expression across 3 biological replicates. Statistics performed with Kruskal Wallis test with Dunn’s multiple comparisons test compared to Halo enrichment. (*) P < 0.05 (****) P < 0.0001. (D) Quantification of integrated FRET density (product of FRET+ percentage and median FRET intensity) of top 10% Halo expressing cells normalized to Halo. N = 6 biological replicates. Statistics performed with Kruskal-Wallis test compared to Halo. (*) P < 0.05. (E) Representative images of Halo/ Polyserine_42_-Halo/Polyserine_42_-Halo variants (magenta) expressed in Wild Type HEK293T cells for 48 hours, and fixed, nuclei stained with DAPI (blue). (F) Quantification of percent cells expressing Halo/ Polyserine_42_-Halo/ Polyserine_42_-Halo variants with at least one Halo/ Polyserine_42_-Halo/Polyserine_42_-Halo variant assembly. N = 15 images across 3 biological replicates. Statistics performed with Kruskal-Wallis test. (****) P < 0.0001.

We began by examining the ability of polyserine_42_ variants to localize to tau aggregates in cells following transfection. First, we observed the replacement of some serine residues with aspartic acids did not alter recruitment into tau aggregates, as constructs with 50% or 25% of the serine residues replaced by aspartic acids (S3D3)7, or (S3D)10SD, respectively, still enriched with tau aggregates (Fig 5B and C). In contrast, polyserine domains with 50% substitution with lysine (S3K3)7, or a threonine rich variant, (ST5)7, failed to colocalize with tau aggregates (Fig 5B and C).

The observation that negatively charged amino acids could substitute serine raised the possibility serine phosphorylation may mediate polyserine interactions with tau aggregates in cells. However, three observations suggest this is not the case. First, the substitution of serine with threonine ((ST5)7), which are often substrates for shared serine/threonine kinases, led to a construct that abolished enrichment in tau aggregates (Fig 5B and C). Second, phos-tag analysis (18) of overexpressed polyserine_42_ did not show a substantial shift in molecular weight, indicating no phosphorylation (Figure S6A). Finally, we note that recombinant polyserine_42_, which is not phosphorylated, can interact with tau *in vitro* (Figure 3) demonstrating phosphorylation is not required for polyserine_42_-tau interaction.

We also examined how these variants altered tau aggregation in cells using flow cytometry (7, 15). We observed that one of these variants retained the ability to increase tau fibrillization. Specifically, we observed that, like polyserine_42_, (S3D)10SD increased tau aggregation (Fig 5D). In contrast, (ST5)7, and (S3K3)7, which do not enrich in tau aggregates, also do not increase tau aggregation (Fig 5B-E). Interestingly, (S3D3)7, which enriches with tau aggregates, does not increase tau aggregation (Figure 5B-D). This demonstrates that the interaction of polyserine domains with tau aggregates can be separated from their ability to promote tau aggregation.

Interestingly, we observed a correlation between the ability for a variant to form assemblies with its ability to increase tau aggregation. Specifically, we observed that polyserine_42_ and (S3D)10SD, both of which increase tau aggregation, formed intracellular assemblies (Figure 5E and F). In contrast, (ST5)7, (S3K3)7, and (S3D3)7 all failed to form intracellular assemblies and to promote tau aggregation (Fig 5E and F).

### Polyserine_42_ Self-Assembly Stimulates Tau Aggregation

To more directly test the role of polyserine_42_ assembly driving the stimulation of tau aggregation, we overexpressed a maltose binding protein (MBP)-polyserine_42_-Halo construct in tau biosensor cells (15) to reduce polyserine_42_ assembly, as MBP has been used to increase solubility of recombinant proteins for structural biology (19).

First, we ensured that MBP fusion reduced polyserine_42_ assembly but retained enrichment in tau aggregates. We found that MBP-polyserine_42_-halo expression reduced the fraction of cells with assemblies to levels similar to MBP-Halo expression alone (Fig 6A and B). Second, we confirmed that MBP-polyserine_42_ enriches with tau aggregates (Fig 6C and D).

**Figure 6:**
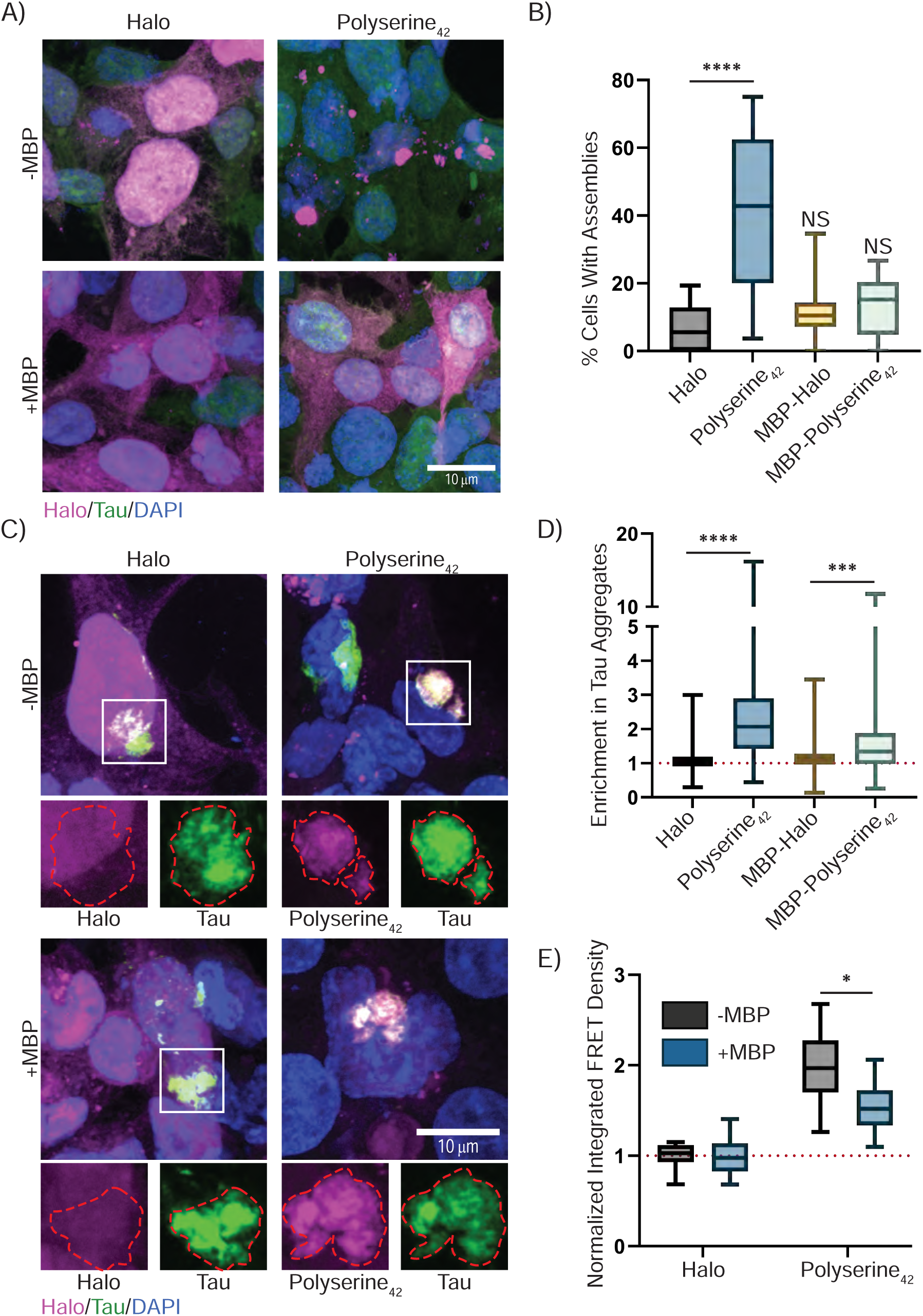
MBP-Polyserine_42_ Fusion Protein Blocks Polyserine Assembly and Stimulation of Tau Aggregation. (A) Representative images of HEK293T tau biosensor cells transfected with plasmids expressing Halo, Polyserine_42_-Halo, MBP-Halo, or MBP-Polyserine_42_-Halo labeled with JF646 (magenta) for 48 hours, tau-YFP (green), and nuclei stained with DAPI (blue). (B) Quantification of percentage of cells expressing Halo tagged constructs with Halo assemblies. N = 15 images across 3 biological replicates. Statistics performed with Kruskal-Wallis test compared to Halo. (****) P < 0.0001 (C) Representative images of HEK293T tau biosensor cells transfected with plasmids expressing Halo, Polyserine_42_-Halo, MBP-Halo, or MBP-Polyserine_42_-Halo labeled with JF646 (magenta) for 24 hours, transfected with tau brain homogenate from aged Tg2541 mice to induce endogenous tau aggregation (green), cells were fixed 24 hours later and nuclei stained with DAPI (blue). Channel breakouts for Halo, Polyserine_42_-Halo, MBP-Halo, or MBP-Polyserine_42_-Halo and tau shown below with tau aggregate perimeter in red superimposed across channels. (D) Quantification of enrichment of Halo, Polyserine_42_-Halo, MBP-Halo, or MBP-Polyserine_42_-Halo in tau aggregates as in (C). N = at least 109 cells with Halo, Polyserine_42_-Halo, MBP-Halo, or MBP-Polyserine_42_-Halo expression and tau aggregates across 3 biological replicates. Statistics performed with Kruskal-Wallis test. (***) P = 0.0001 (****) P < 0.0001.

We next tested if this separation of functions altered tau aggregation. We found that addition of MBP reduced polyserine_42_ effects on promoting tau aggregation (Fig 6D). Therefore, polyserine_42_ self-assembly positively correlates with increased tau aggregation, and that self-assembly and targeting to tau aggregates are separable functions.

## Discussion

In this study we demonstrate that polyserine_42_ spontaneously self assembles. The key observation is that purified polyserine spontaneously forms microscopically visible assemblies large enough to pellet via ultracentrifugation (Fig 2). While the specific structure of these polyserine assemblies is unknown, it may form complex coiled-coil structures similar to serine repeats of shorter lengths (20). Additionally, the self-assembly of polyserine provides a molecular understanding of polyserine assemblies that form upon overexpression in cells and mice (7, 11). Polyserine assemblies are also observed in spinocerebellar ataxia type 8 (SCA8) and Huntington’s Disease (HD), produced by RAN translation, and may form through similar self-assembly mechanisms (21, 22). An interesting possibility is that polyserine domain self-assembly may promote the assembly of nuclear speckles, MIGs, or CSs, as SRRM2 is proposed to contribute to nuclear speckle formation (7, 23).

Our data supports a hypothesis that polyserine assembly is required to stimulate tau aggregation. Importantly, blocking polyserine assembly via expression of an MBP-polyserine fusion protein also reduces stimulation of tau aggregation (Fig 6). Taken together with our findings that polyserine variants which do not assemble also do not increase tau aggregation (Fig 5), we suggest that assembly of polyserine domains stimulates tau aggregation. This may also explain the observation that polyserine puncta and tau aggregation are observed in some cases of SCA8 and HD (24, 25). Further study is required to understand if pathological polyserine assembly directly induces tau aggregation in these diseases.

Polyserine assemblies increase tau aggregation which can occur through direct interaction with tau. We previously observed that overexpression of polyserine_42_ increases tau aggregation in HEK293T tau biosensor cells and pathology in PS19 mice (7, 11). Here, we provide evidence that this increased aggregation can occur due to direct interaction between polyserine assemblies, tau monomers and other cofactors sufficient to induce tau aggregation (Fig 3). Additionally, polyserine containing assemblies can localize tau seeds, creating preferred sites for tau aggregation (Fig 1). Thus, polyserine expression and assembly may exacerbate tau pathology by creating a complex network with tau seeds which promotes subsequent aggregation.

Polyserine assemblies may interact with tau and increase tau aggregation in multiple ways. In one manner, polyserine assemblies enrich tau monomers and cofactors, such as RNA, sufficient for fibrilization in a high local concentration (Fig 3). This higher concentration of tau and RNA may then lead to faster fiber formation and growth (Fig 4). Alternatively, polyserine may interact with tau monomers and bias the folding of tau into an aggregate prone form. Finally, polyserine assemblies may increase tau aggregation by enriching seeding competent oligomers or fibers with tau monomers, thus increasing the rate of fiber growth independent of additional cofactors.

In conclusion, we hypothesize that polyserine domains self-assemble, which increases the local concentration of tau monomers and fibers through direct interactions, resulting in increased rates of tau aggregate formation (Fig 7).

**Figure 7:**
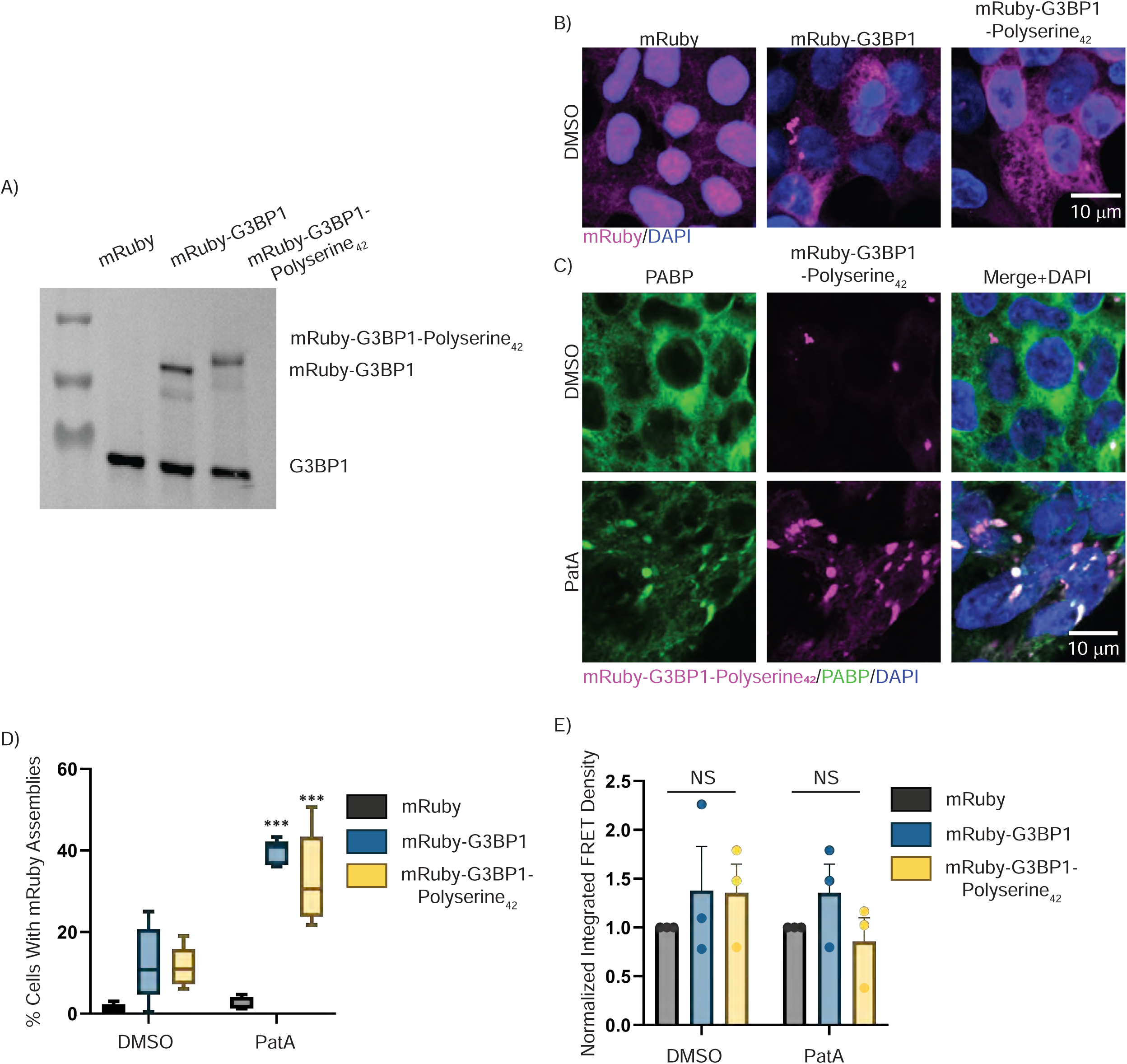
Polyserine_42_ Assemblies are Preferred Sites of Tau Aggregation Through Enrichment of Tau Seeds Stimulating Tau Aggregation. (A) Model showing a possible mechanism in which polyserine containing assemblies enrich tau monomers, RNA, and tau seeds to create preferred sites of tau aggregation, which in turn stimulates tau aggregation. Tau seeds can enrich in cytoplasmic polyserine containing assemblies where they interact with endogenous tau, and possibly other cellular cofactors to grow a tau aggregate.

## Experimental procedures

### Cell culture and tau aggregate seeding of HEK293T cells

HEK293T tau biosensor cells(15) stably expressing the 4R repeat domain of tau (K18) with the P301S mutation were purchased from ATCC (CRL-3275). Cells were plated at 1.25 X 10^5^ cells/mL in 500 uL of DMEM supplemented with 10% FBS and 0.2% penicillin-streptomycin on poly-D-lysine coated glass coverslips for immunofluorescence and microscopy, or directly in a 24-well cell culture treated plate (Corning 2536) for flow cytometry. Cells were allowed to grow overnight at 37 °C and 5% CO2. The next day, plasmids were transfected using lipofectamine 3000 following manufacturers protocol. For cells expressing Halo tagged constructs, Janelia Fluor 646 (Promega) was added for downstream imaging, or TMRDirect Ligand (Promega) was added for downstream flow cytometry. The next day, cells were transfected with lipofectamine 3000 with 0.25 uL of 1 mg/mL clarified brain homogenate from Tg2541 mice per well. 24 hours later, cells were either fixed for immunofluorescence and imaging or were processed for flow cytometry (see below).

### Clarification of brain homogenate for tau aggregate seeding in HEK293T tau biosensor cells

As previously described (6), 10% brain homogenate from Tg2541 mice was centrifuged at 500 x g for 5 minutes. The supernatant was collected and centrifuged again at 1000 x g for 5 minutes. The supernatant was collected, protein concentration determined with a bicinchoninic acid assay (BCA) and diluted with PBS to 1 mg/mL. Aliquots were stored at -80C for downstream transfection.

### Generation of cell lines

For production of mRuby-G3BP1-polyserine_42_ expressing HEK293T tau biosensor lines, construct encoding mRuby-G3BP1-polyserine_42_ was cloned into a pLenti EF1 vector. Wildtype HEK293T cells were transfected using Lipofectamine 3000 with pLenti plasmids and lentiviral packaging plasmids (Gag-pol (Addgene #14887), VSV-G (Addgene #8454), rSV-REV (Addgene #12253). Lentivirus was collected 48 hours later and was added to HEK293T tau biosensor cells described previously (15). Transduced cells were selected with blasticidin (2 µg/mL).

### Immunofluorescence and Fixed Cell Imaging

As described above, cells plated and prepared on poly-D-lysine coated coverslips were fixed with 4% paraformaldehyde for 15 minutes. Cells were washed once with 1xPBS, permeabilized with 0.5% TritonX for 10 minutes, washed three times with 1x PBS, and blocked with 5% BSA in PBS for 1 hour at room temperature. Primary antibodies were added in blocking buffer for two hours at room temperature (Rabbit anti-PolyA Binding Protein (abcam 21060 1:500 in TBS-T)). Cells were washed three times with 1x PBS. Secondary antibody (Anti-Rabbit-AlexaFluor 549 (abcam150092 1:1000 in TBS-T)) solution in blocking buffer with 1x DAPI was added to cells for 1 hour. Cells were washed three times with 1x PBS and mounted using ProLong Glass Antifade Mountant.

For monitoring enrichment of stress granules with tau aggregates, cells were plated as described above. Twenty-four hours after plating, cells were transfected with tau brain homogenate (described above) and seven hours later treated with 50 nM PatA. Twelve hours later, cells were fixed and mounted as described above.

For monitoring enrichment of polyserine and polyserine variants with tau aggregates, cells were plated as described above. Twenty-four hours after plating, cells were transfected with plasmids encoding Halo tagged constructs alongside JF646 Halo-ligand (Promega). Twenty-four hours after transfection, tau aggregates were induced as described above. Twenty-four hours after tau brain homogenate transfection, cells were fixed and mounted as described above.

### Live Cell Imaging

For analysis of tau aggregation in the context of stress granules with and without polyserine, cells were plated on 24 well poly-D-lysine glass bottom plates with #1.5 cover glass at 0.2 x 10^6^ cells/mL. Cells were allowed to grow overnight at 37 °C and 5% CO2. Tau aggregation was induced 24 hours later with brain homogenate from Tg2541 mice as described above, seven hours later, stress granules were induced with 50 nM PatA and nuclei were stained with Hoechst. Images were collected every 10 minutes for 16 hours.

For live cell imaging of tau seeds, cells were plated as described above. For localization with SRRM2 or PNN, JF549 was added alongside initial plating. Cells were allowed to grow overnight at 37 °C and 5% CO2. Fluorescent tau fibers were prepared (see below) pelleted and 100K x g for 30 minutes, washed once in 1xPBS, pelleted again at 100K x g for 30 minutes and resuspended in 1xPBS. Tau fiber pellets were then sonicated just prior to transfection (performed identically to transfection of tau brain homogenate from Tg2541 mice). For localization to stress granules, cells were treated with 50 nM PatA 7 hours after tau fiber transfection alongside nuclei staining with Hoechst just prior to imaging. For localization to SRRM2 or PNN assemblies, tau fiber transfection occurred alongside nuclei staining with Hoechst just prior to imaging. Images were collected every 10 minutes for 16 hours.

### Cloning of polyserine expression plasmids

To express polyserine variants in HEK293T tau biosensor cells(15) custom gBlocks were ordered from Twist Bioscience and cloned into pCDNA3.1-Halo plasmids under a CMV promoter using In-Fusion cloning. Bacterial expression plasmids were produced using custom gBlocks ordered from Twist Bioscience or IDT which were cloned into pET28 vectors using In-Fusion cloning.

### Protein expression and purification

pET28 plasmids expressing SUMO-Polyserine_42_-Halo, SUMO-Halo, SUMO-Polyserine_42_-Cys, SUMO-Cys, and pET29 2N4R P301S single cysteine (See Fig 3A) were transformed into Rosetta2(DE3)pLysS *E. coli* and grown in LB media supplemented with kanamycin and chloramphenicol. One liter LB cultures were inoculated with overnight cultures, incubated with shaking at 37 °C until O.D600 >= 0.4, and protein expression was induced with 0.2 mM IPTG (polyserine constructs), or 0.5 mM IPTG (Halo and SUMO-Cys constructs), or 0.4 mM IPTG (tau constructs) for 4 hours at 37°C. Bacteria was pelleted and stored at -80 °C.

All SUMO tagged constructs were purified using Ni-NTA columns. Cell pellets were resuspended in Lysis Buffer (50 mM MOPS pH 7.0, 300 mM NaCl, 30 mM imidazole, 1 mM DTT, supplemented with Ultra EDTA-free protease inhibitors and 0.1 mM AEBSF) and lysed with sonication. Cell lysate was clarified with centrifugation and filtered through a 0.2 µm pore filter. For downstream imaging with halo-ligands, JF594 or JF646 was added to clarified lysate and nutated at room temperature for 30 minutes. Labeled protein lysate was added to equilibrated Ni-NTA, washed with 5 column volumes (CV) of lysis buffer, 10 CV of lysis buffer with 500 mM NaCl, and finally 10 CV of lysis buffer with 1M NaCl. Six CV of Elution buffer (50 mM MOPS pH 7.0, 200 mM NaCl, 300 mM imidazole, 1 mM DTT, supplemented with Ultra EDTA-free protease inhibitor (Sigma) and 0.1 mM AEBSF) was added and incubated with resin for 30 minutes. Elution buffer was collected and concentrated before overnight dialysis into storage buffer (50 mM MOPS pH 7.0, 100 mM NaCl, 1 mM DTT). For maleimide labeling, proteins were purified as described above, but with 1 mM TCEP instead of DTT in all buffers. For maleimide labeling, concentrated, purified protein was incubated with 3 molar excess of maleimide dye for 2 hours at room temperature. Excess dye was removed with overnight dialysis into storage buffer as described above.

Full length tau constructs were purified over three columns, Ni-NTA, SP cation exchange, and S200 size exclusion. Clarified cell lysates were prepared as described above in Tau Lysis Buffer (50 mM Tris pH 7.4, 300 mM NaCl, 30 mM imidazole, 1 mM DTT, supplemented with Ultra EDTA-free protease inhibitor (Sigma)). Lysate was added to equilibrated Ni-NTA, washed with 5 CV of Tau Lysis Buffer, 10 CV Tau wash buffer supplemented with 500 mM NaCl, 10 CV Tau wash buffer with 1 M NaCl, and eluted in Tau elution buffer (50 mM Tris pH 7.4, 100 mM NaCl, 300 mM imidazole, 1 mM DTT). Eluted protein was dialyzed overnight into Buffer A (50 mM MES pH 6.00, 150 mM NaCl, 1 mM DTT). Protein was loaded into a superloop on an AKTA Pure 25 M and passed through a pre-equilibrated SP cation exchange column. Protein was eluted with a gradient of Buffer B (Buffer A with 1 M NaCl) and fractions collected. Protein containing fractions were pooled, concentrated to 1 mL and loaded onto an S200 size exclusion column. Protein was eluted in storage buffer (1x PBS pH 7.4, 1 mM TCEP (for maleimide labeling) or 1 mM DTT (for fibrilization reactions). Protein containing fractions were pooled and concentrated to 50 µM and stored at 4 °C.

### Microscopy Colocalization

Colocalization assays were performed in 1xPBS with 3 µM polyserine or halo proteins and 5 µM of experimental proteins. Protein mixture was brought up to 50 µL in Bio-One CELLview slides (Greiner) and immediately imaged. Imaging was performed on a Nikon spinning disk confocal microscope at 60x magnification.

### *In vitro* polyserine assembly

To assess assembly formation, 20 µM purified SUMO-Halo and SUMO-polyserine_42_-Halo were incubated with TEV for 24 hours at room temperature in storage buffer. Reactions were either subjected to ultracentrifugation for dot blots, or microscopic examination. For ultracentrifugation analysis, reactions were pelleted at 100K x g for 1 hour, supernatant kept, and pellet washed with dialysis buffer, and pelleted again at 100K x g for 30 minutes. 1 µL of total, supernatant, and pellet fractions were spotted onto nitrocellulose and blotted with anti-HaloTag pAb (Promega). For microscopic examination, reactions were transferred to Bio-One CELLview slides (Greiner) and imaged at 60x magnification on a Nikon spinning disk confocal microscope.

### Preparation of Recombinant Tau Fibers

Tau fibers were prepared using JF646-maleimide labeled tau monomers purified and stored in TCEP as described above. For fibrilization, 5 µM tau monomers were incubated with 40 ng polyU RNA in 1xPBS pH 7.4 and freshly prepared 1 mM DTT for 24 hours with shaking at 37 °C. Reactions were done in separate 50 µL reactions and pooled after incubation. Tau fibers were separated from soluble tau species via ultracentrifugation. Pooled tau fiber reactions were centrifuged at 100K xg for 30 minutes, the supernatant removed, and the pellet resuspended in equal volumes of 1x PBS pH 7.4. The resuspended pellet was centrifuged at 100K xg for 30 minutes, the supernatant removed, and the pellet resuspended in 1x PBS pH 7.4. The resuspended pellet was sonicated and immediately used in downstream experiments.

### Preparation of Fluorescent RNA

Fluorescent polyU RNA homopolymers were prepared using poly-uridylic acid (Millipore Sigma) and the Label IT® Nucleic Acid Fluorescein Labeling Kit (Mirus Bio) according to manufacturers protocol. Fluorescent RNA was labeled on the day of recombinant tau fiber reactions.

### Thioflavin-T kinetics assays

To assess fibrilization kinetics, 5 µM unlabeled 2N4R tau purified and stored in 1 mM DTT was mixed with 40 ng/uL polyU RNA, 3 µM SUMO-polyserine_42_-Halo or SUMO-Halo, and 20 µM thioflavin T in 1x PBS pH 7.4 supplemented with freshly prepared 1 mM DTT. Reactions were prepared in 175 uL master mix and 50 µl was aliquoted into three wells of a 96 well #1.5 cover glass bottom plate (Greiner). Negative controls without tau, with and without RNA, and with and without additional proteins were also prepared in triplicate. Thioflavin T fluorescence was measured every 5 minutes for 24 hours on a BMG CLARIOstar plate reader using 446 nm excitation and 482 nm emission, incubating at 37°C and shaking at 300 RPM for 30 seconds prior to each measurement. Resulting curves from 3 technical replicates and 3 biological replicates, for 9 total replicates, were plotted and globally fit to logistic growth curve using Prism 10.4.2.

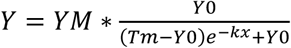

The rate constant (k) and lag time (1/k, or time of first inflection point) were extracted from the fit equation and plotted with SEM from 9 technical replicates.

### Immunogold labeling and TEM

For TEM, tau fibers were made by incubating 5 µM monomeric 2N4R P301S Tau, 40 ng PolyU RNA, 1 mM DTT, with and without 3 µM SUMO-polyserine_42_-Halo or SUMO-Halo, in 1x PBS pH 7.4 for 24 hours at 37°C while shaking at 300 RPM. Tau fibrilization samples were diluted 1:5 and 5 µl of each were adsorbed on a carbon-coated copper grids (300 mesh, Electron Microscopy Sciences) for 1 min. Excess sample was wicked off and grids were incubated with 5 µl 2% uranyl-acetate (UA) for 30 seconds. Excess UA was wicked off and grids were allowed to dry.

For immunogold labeling, 10 µl of diluted tau fibrilization samples (See above) were adsorbed on carbon-coated nickel grids (300 mesh, Electron Microscopy Sciences) for 10 minutes. Excess fibrils were removed by placing the grids on three successive drops of fresh 1x PBS without drying. Grids were blocked in 1% BSA in PBS for 60 min and subsequently incubated with primary anti-Halo antibody (anti-HaloTag pAb (Promega)) diluted 1:100 in 1% BSA in PBS) for 60 min. The grids were washed on five successive drops of fresh 1x PBS without drying and incubated with gold-labeled secondary antibody (10 nm-Gold goat anti-rabbit IGG (Ted Pella) diluted 1:50 in 1% BSA) for 60 min. Grids were washed again on five successive drops of fresh 1x PBS and samples were negatively stained by incubating in 5 µl 2% UA for 30 seconds. TEM was performed with a FEI Tecnai T12 spirit equipped with an AMT CCD camera.

### SDS-PAGE and Western Blotting

For analysis of proper expression of mRuby-G3BP1 and mRuby-G3BP1-Polyserine_42_, cells were grown in 6 well plates to 80% confluence. Cells were harvested in 1x PBS and pelleted at 300 xg for 8 minutes. The cell pellet was resuspended in lysis buffer (10 mM Tris pH 7.4, 2.5 mM MgCl2, 100 mM NaCl supplemented with PhosStop (Sigma) and Ultra Complete Protease Inhibitor (Sigma)). Cells were lysed via pipetting through a 26g needle 10 times. Lysate was clarified via centrifugation at 10K xg for 15 minutes at 4 °C. Supernatant was collected and stored at -80.

Cell lysate protein concentration was measured with Bradford assay and normalized to the lowest protein concentration. Lysate mixed with 4X SDS-Loading dye and boiled for 5 minutes prior to loading on a NuPAGE 4-12% Bis-Tris mini gel (Invitrogen). Protein was transferred to nitrocellulose membrane which was then blocked with 5% BSA in TBS-T. The blocked membrane was incubated with primary antibody solution diluted in 5% BSA in TBS-T overnight at room temperature, washed 3 times with TBS-T, and incubated with secondary antibody solution for one hour followed by 3 washes with TBS-T. Blot was developed with Clarity Western ECL Substrate (BioRad).

For Phos-Tag analysis of polyserine_42_ from HEK293T cells, cells were grown in 6 well plates to 50% confluence. Cells were transfected with plasmids expressing polyserine_42_-halo or halo and incubated at 37 °C and 5% CO2 for 48 hours. Cells were harvested in 1x PBS and pelleted at 300xg for 8 minutes. The cell pellet was resuspended in lysis buffer (10 mM Tris pH 7.4, 2.5 mM MgCl2, 100 mM NaCl, supplemented with Ultra Complete Protease Inhibitor (Sigma)). Resuspended cell pellets were split into two fractions and diluted with either lysis buffer with or without PhosStop (Sigma). Cells were then lysed via pipetting through a 26g needle 10 times. Lysate was clarified via centrifugation at 10K xg for 15 minutes at 4 °C. Supernatant was collected and total protein concentration was measured with Bradford assay and normalized to the lowest protein concentration. Lysates were then treated with lambda protein phosphatase or buffer according to manufacturer protocol (New England Biolabs). Treated supernatants were then mized with 4X SDS-Loading dye and boiled for 5 minutes prior to loading on SuperSep™ Phos-tag™ gels (FUJIFILM Wake Pure Chemical Corporation). The gel was transferred to a nitrocellulose membrane after incubation in 10 mM EDTA as described above. Western blot was performed as described above.

### Flow Cytometry

Flow cytometry was used to determine the fraction of cells with tau aggregates as described previously(7). Briefly, HEK293T tau biosensor cells(15) were plated at 0.125 x 10^6^ cells/mL in 500 µL in a 24 well plate. For halo expression, 24 hours after plating, plasmids expressing halo tagged constructs were transfected with lipofectamine 3000 following manufacturers protocol. At the time of transfection, 200 nM of TMRDirect halo-ligand was added. Twenty-four hours after transfection, tau aggregates were induced as described above using 0.5 ng/µL of tau brain homogenate. Twenty-four hours after transfection of tau brain homogenate, cells were trypsinized, washed with PBS and filtered with 50 µm nylon mesh filters prior to analysis. Sorting was performed on BD FACSCelesta™ Cell Analyzer with the following filter sets: 561-585 (Halo expression), 405-450 (CFP), and 405-525 (FRET). Analysis was performed using FlowJo. Gating was performed sequentially, first gating for cells, single cells, then Halo expression for top 10% expressing cells, and finally, gated for FRET+ cells. The FRET+ gate was set using a mock transfected well with 1% as previously detailed(15). The integrated FRET density was determined as a product of the median FRET intensity and the percentage of FRET+ cells.

For analysis of expression of G3BP1 and G3BP1-Polyserine_42_ with and without stress, cells were plated at the same density. Cells were transfected with tau brain homogenate 24 hours after plating as described above. Seven hours after transfection of tau brain homogenate, cells were treated with 50 nM PatA. Twelve hours after PatA treatment, cells were then processed and analyzed as described above.

### Quantification and statistical analysis

Statistical details of experiments are included in the figure captions (including statistical tests used, value of n, what n represents). All error bars reported are SEM. All statistical analyses were performed using GraphPad-Prism 10.

### Image Analysis

All image analysis was performed with Cell Profiler version 4.2.8.

To measure the fraction of cells with mRuby or Halo positive assemblies, as well as the enrichment of mRuby or Halo fluorescence in tau aggregates, the same pipeline was used. Briefly, cell nuclei and cytoplasm were segmented and filtered for Halo or mRuby expressing cells. This population of cells was then further filtered for cells with and without tau aggregates, or with and without mRuby or Halo assemblies. For enrichment of Halo or mRuby, the mean intensity was determined inside tau aggregates, and from the remainder of the cytoplasm on a single cell basis. Enrichment was determined as the mean intensity of Halo or mRuby inside tau aggregates divided by the mean intensity of Halo or mRuby in the remainder of the cytoplasm.

To determine the enrichment of proteins in polyserine assemblies, fields of view were segmented for polyserine assemblies, these assemblies were then merged as one object per field of view. The mean intensity of the additional proteins was then measured in and outside polyserine assemblies, and enrichment was determined with the same method described above. This pipeline was also used to measure the area of polyserine assemblies on a per assembly basis.

## Data availability

### Lead Contact

Further information and requests for resources and reagents should be directed to and will be fulfilled by the lead contact, Roy Parker.

### Materials availability

All unique/stable reagents generated in this study are available from the Lead Contact without restriction.

### Data and code availability

Microscopy data reported in this paper will be shared by the lead contact upon request. Any additional information required to reanalyze the data reported in this paper is available from the lead contact upon request.

## Acknowledgements

Electron microscopy was done at the University of Colorado, Boulder EM Services Core Facility in the MCDB Department, with the technical assistance of facility staff. We would also like to thank Meaghan Van Alstyne and Carolyn Decker for their input on experimental design and proofing the text and figures.

## Author Contributions

J.P, K.M, and R.P designed the study. J.P., K.M and J.K performed experiments. J.P, K.M and R.P wrote and revised the manuscript.

## Funding and additional information

This work was supported by funds to R.P from the Howard Hughes Medical Institute. Flow cytometry was performed in University of Colorado Department of Biochemistry flow cytometry core with funding from SCR_019309, with instruments purchased with funding provided by grant S10OD021601. Cell culture was performed at the University of Colorado Department of Biochemistry cell culture core facility with funding from SCR_018988.

---

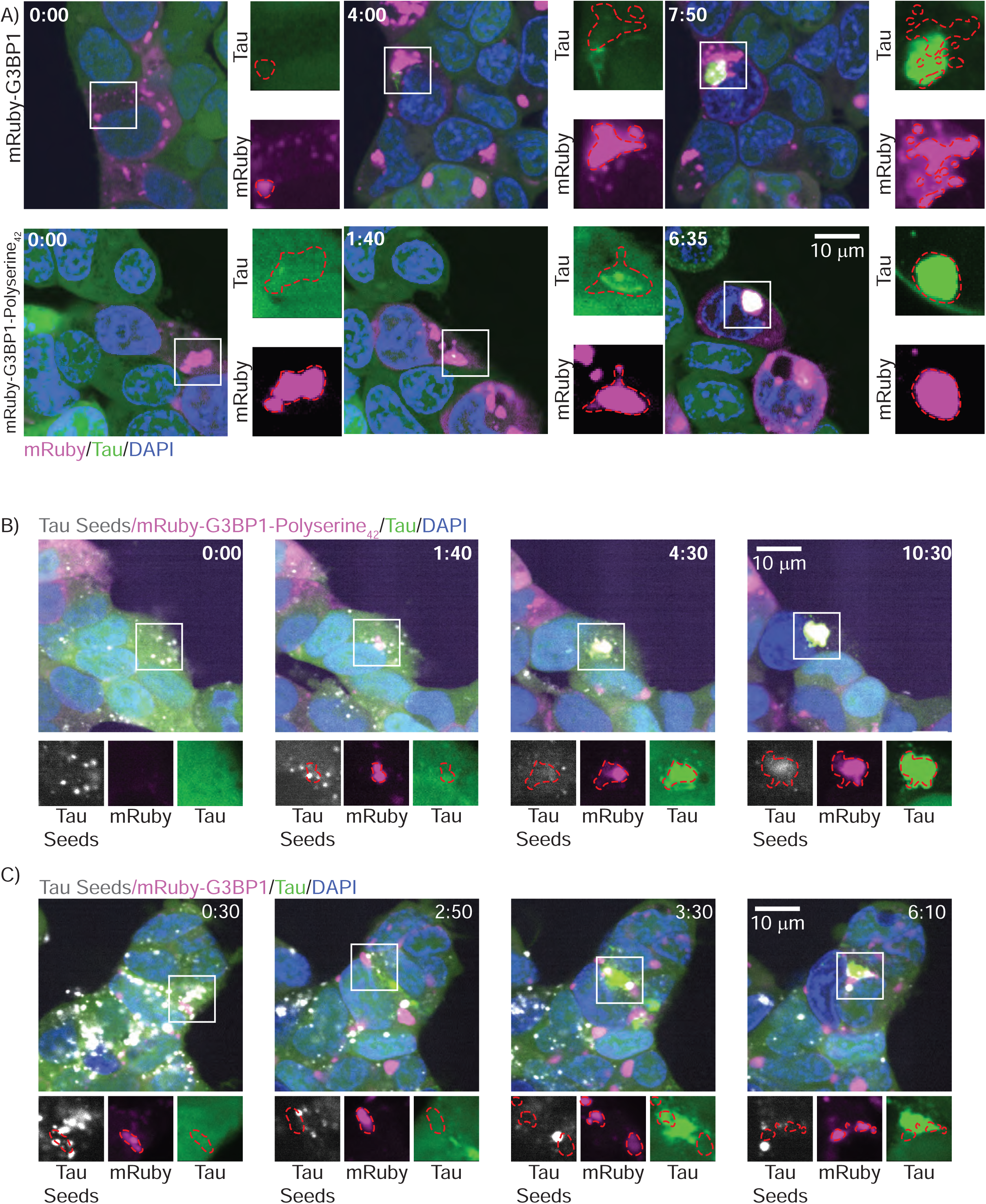

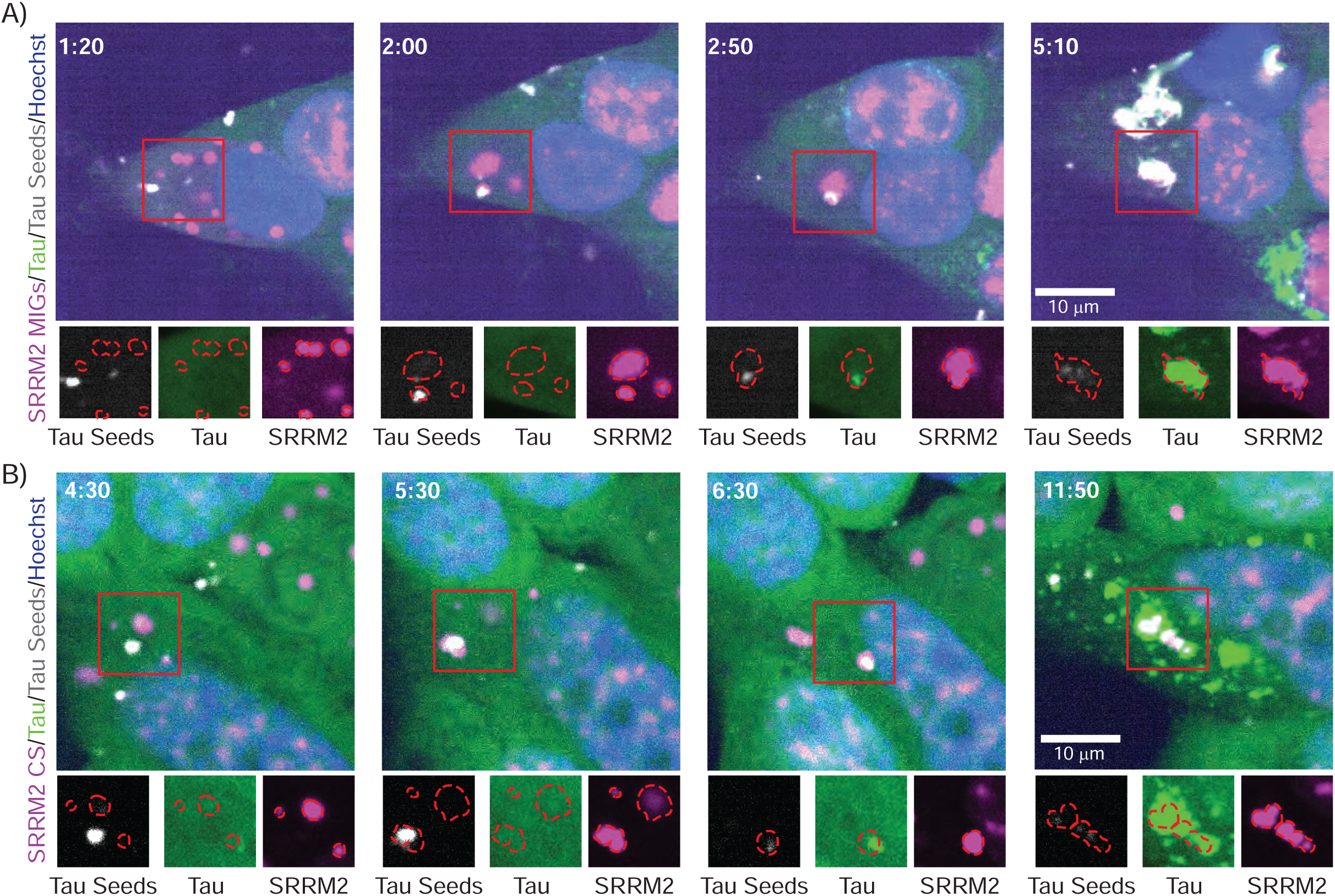

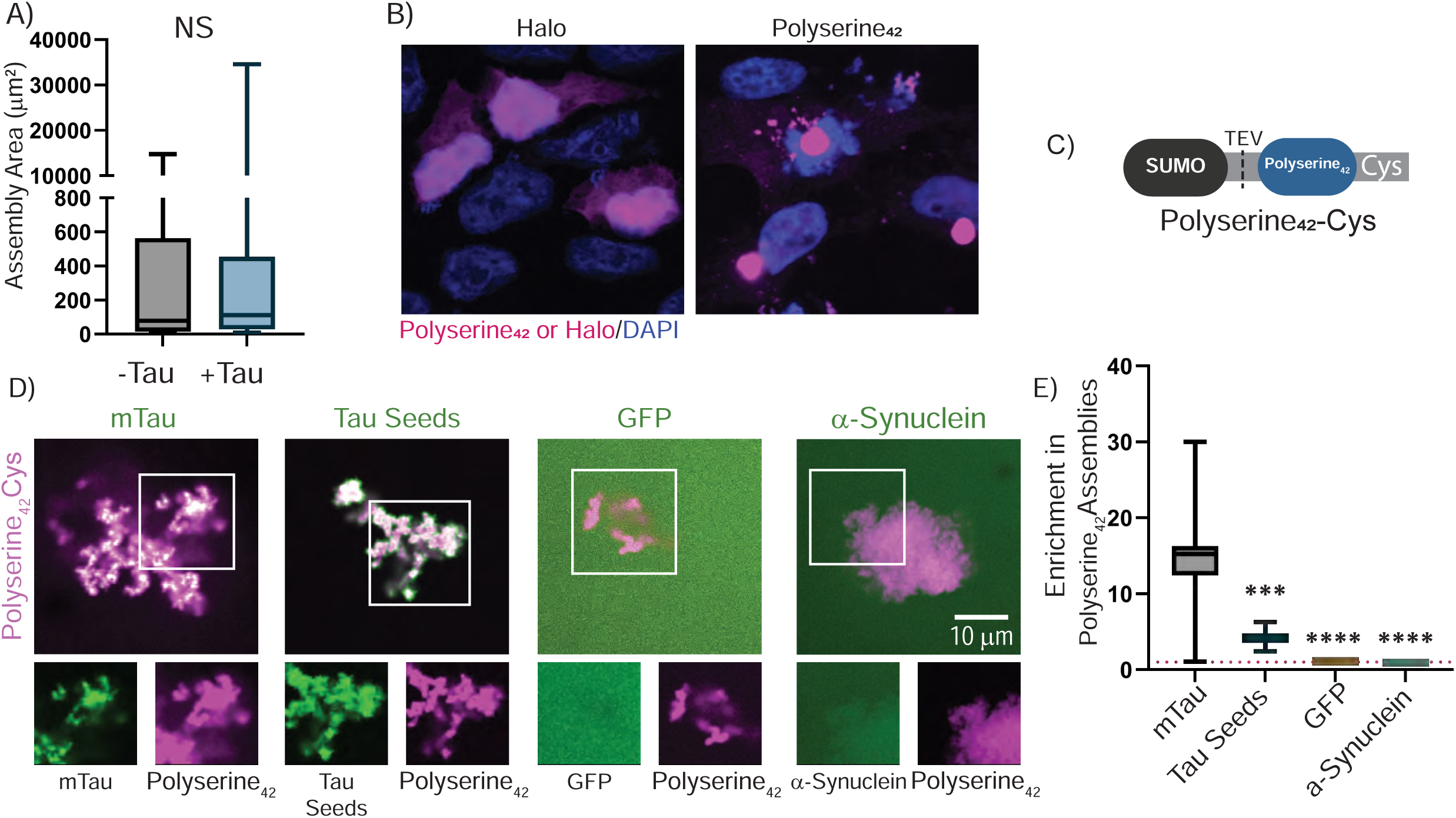

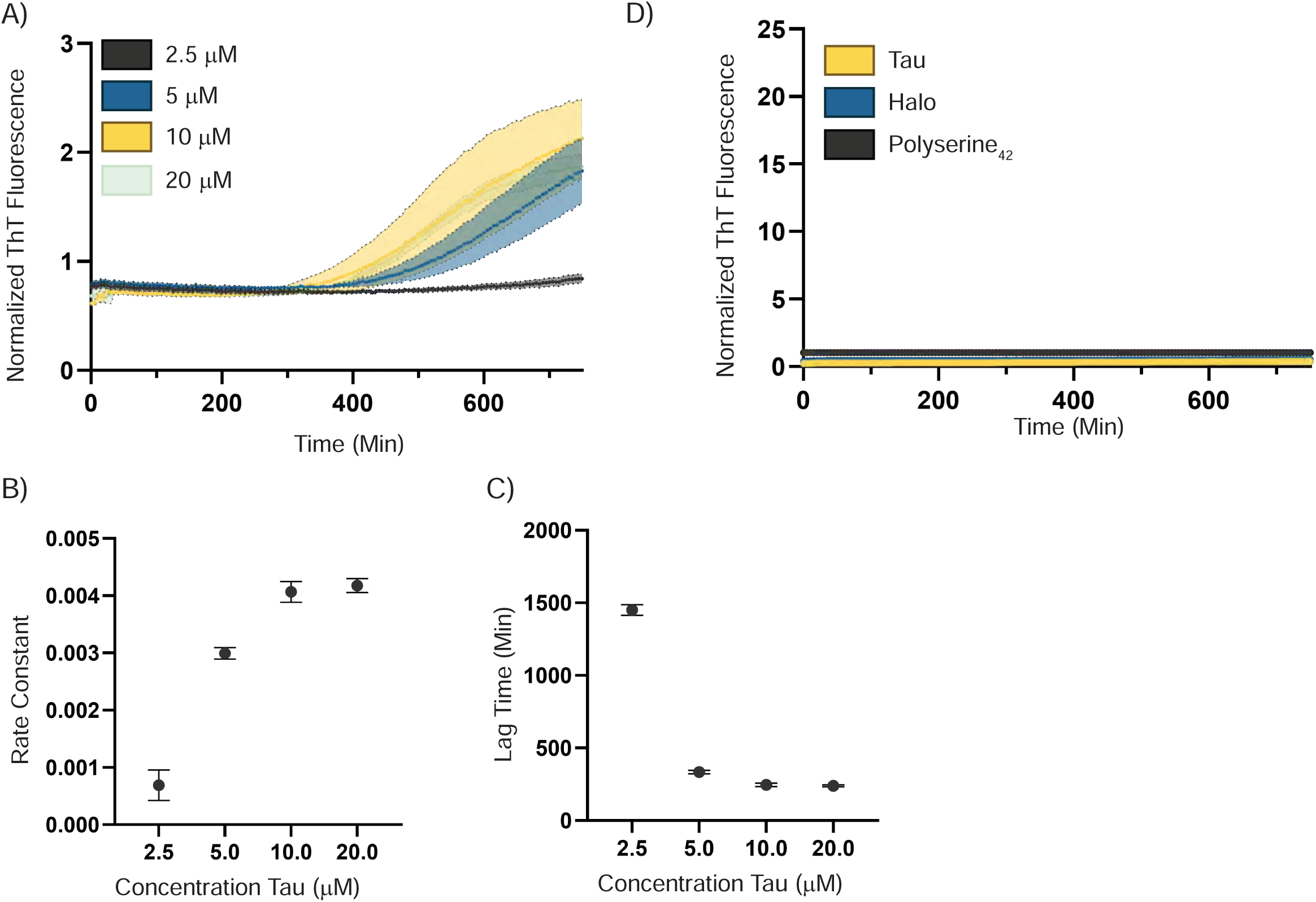

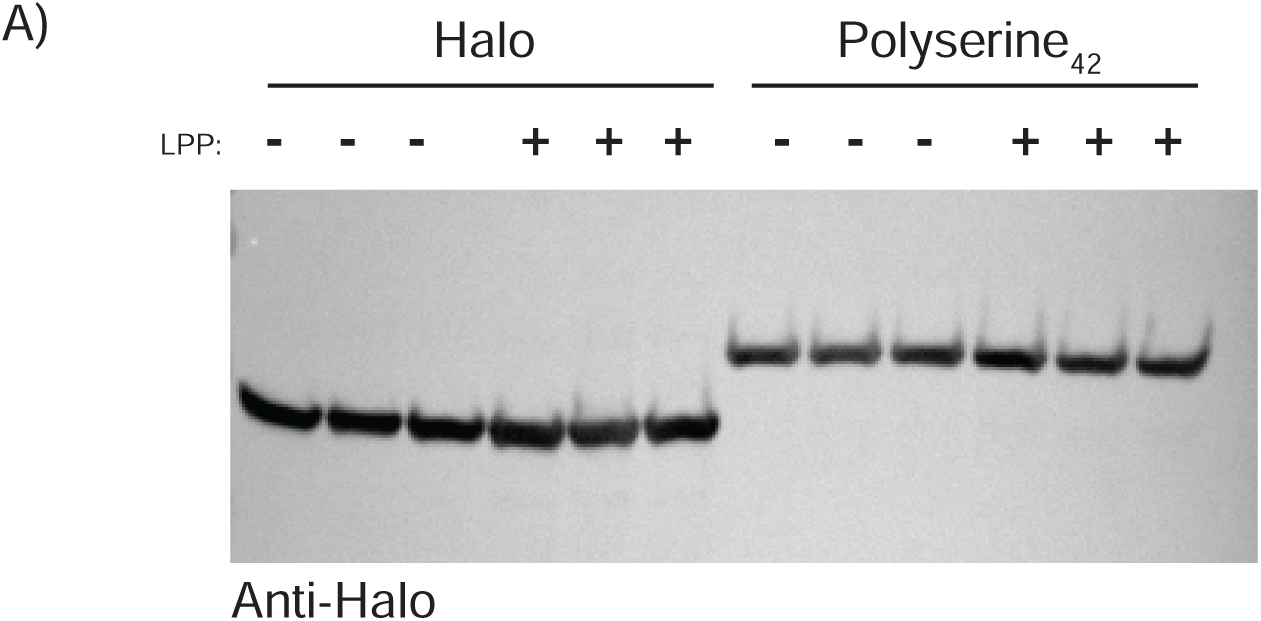

